# Longitudinal analysis of the humoral response to SARS-CoV-2 spike RBD in convalescent plasma donors

**DOI:** 10.1101/2020.07.16.206847

**Authors:** Josée Perreault, Tony Tremblay, Marie-Josée Fournier, Mathieu Drouin, Guillaume Beaudoin-Bussières, Jérémie Prévost, Antoine Lewin, Philippe Bégin, Andrés Finzi, Renée Bazin

## Abstract

Héma-Québec, the blood supplier in the Province of Quebec, Canada, collects and tests convalescent plasma used in a clinical trial to determine the clinical efficacy of this product for the treatment of hospitalized COVID-19 patients. So far, we have collected 1159 plasma units from 282 COVID-19 convalescent donors. The presence of antibodies to the receptor binding domain (RBD) of SARS-CoV-2 spike protein in convalescent donors was established at the first donation. Seropositive donors were asked to donate additional plasma units every six days. Until now, 15 donors have donated at least four times and, in some cases, up to nine times. This allowed us to perform a longitudinal analysis of the persistence of SARS-CoV-2 RBD-specific antibodies in these repeat donors, with the first donation occurring 33-77 days after symptoms onset and donations up to 71-114 days after symptoms onset thereafter. In all donors, the level of antibodies remained relatively stable up to about 76 days after symptoms onset but then started to decrease more rapidly to reach, in some convalescent donors, a seronegative status within 100-110 days after symptoms onset. The decline in anti-RBD antibodies was not related to the number of donations but strongly correlated with the numbers of days after symptoms onset (r = 0.821). This suggests that *de novo* secretion of SARS-CoV-2 RBD antibodies by short-lived plasma cells stopped about 2-3 months after disease onset, an observation that has important implications for convalescent plasma collection and seroprevalence studies undertaken several months after the peak of infection.

## MAIN TEXT

The search for therapeutic options to treat severely ill COVID-19 patients has prompted the initiation of many clinical trials, some of which are exploring the transfusion of COVID-19 convalescent plasma (CCP) as a means to reduce the severity of the disease and help resolve the infection more rapidly. Beneficial effects of CCP transfusion in COVID-19 patients have been recently reported, although the studies were not controlled randomized trials or involved only a few patients (1–7). One of the main hypotheses to explain the potential clinical benefits of CCP is the presence of SARS-CoV-2 neutralizing antibodies (nAb) (8,9). Therefore, several groups have included nAb titers as a criterion for the selection of CCP units to be transfused (10,11). However, the determination of nAb titers using a virus neutralization assay is not readily accessible and alternative ways have been considered. In this regard, several reports have shown a very good correlation between nAb and SARS-CoV-2 spike protein receptor binding domain (RBD) antibody titers (11–15). Analysis of SARS-CoV-2 spike RBD antibodies using ELISA is thus a valuable tool for the initial characterization of CCP to be used in clinical trials.

Héma-Québec, the agency responsible for the blood supply in the province of Quebec, Canada, is involved in the collection and testing of CCP used in a clinical trial (CONCOR-1 trial, ClinicalTrials.gov Identifier: NCT04348656) designed to determine the effect of CCP at reducing the risk of intubation or death in adult patients hospitalized for COVID-19 respiratory illness. Potential donors were recruited after at least 14 days of resolution of COVID-19 symptoms. Initial diagnosis had been confirmed by public health authorities through either PCR or epidemiologic contact. All participants also met the donor selection criteria for plasma donation in use at Héma-Québec and have consented to the study. To measure the presence of SARS-CoV-2 antibodies in CCP, we developed an in-house ELISA using recombinant SARS-CoV-2 spike RBD as the target antigen and a polyvalent anti-human Ig-HRP conjugate as secondary antibody allowing the detection of all classes of RBD-specific antibodies. The assay was first used to establish seropositivity, which is mandatory to qualify CCP for participation in the CONCOR-1 trial, and is also currently used for additional characterization of CCP such as anti-RBD titers for each Ig class (G, M, A) and for an ongoing seroprevalence study in blood donors. To establish seropositivity, a plasma dilution of 1:100 was chosen since it was shown to easily distinguish between seronegative and seropositive samples as well as provide a semi-quantitative evaluation of the level of anti-RBD antibodies over a range of optical density (OD_450 nm_) from the negative control value (around 0.070) up to 2.950. The initial cut-off value for seropositivity, set at 0.250, was calculated using the mean OD + 3 standard deviations of 13 COVID-19 negative plasma samples (collected in 2019, before the outbreak of SARS-CoV-2) plus a 15% inter-assay coefficient of variation. Following implementation of the assay, we used results from the analysis of 94 convalescent plasma samples and 66 negative samples and the determined cut-off value yielded a sensitivity of 97.9% and a specificity of 98.5%.

Consistent with previous reports on the rate of seroconversion of COVID-19 patients (16–18), the overall proportion of our convalescent plasma donors (n=282) that were tested seronegative at the time of donation was 6.9%. However, we noted that this proportion increased to about 15% if we considered only donors who had waited for more than 11-12 weeks after symptoms onset before donating (Table 1). This prompted us to perform a longitudinal analysis of the anti-RBD antibody response in CCP donors within a group of 15 individuals (11 males and 4 females, median age of 56 years old, range 20-67) who donated at least four times, during a time interval after symptoms onset ranging from 33-77 days for the first donation to 66-114 days for the last donation (Table 2). These donors reported symptoms of different intensity, ranging from mild/moderate to severe symptoms, although none of them were hospitalized for COVID-19.

**Table 1:** COVID convalescent plasma donors characteristics.

|  | All donors | Male | Female |
| --- | --- | --- | --- |
| Donors (n) | 282 | 158 | 124 |
| Age (mean ± SD) | 37 ± 13 | 42 ± 14 | 31 ± 9 |
| Age (median) | 33 | 39.5 | 28 |
| Seronegative at first donation* | 6.9% | 3.0% | 12.4% |
| Seronegative; first donation 78-109 days<br>after symptoms onset | 15.0% | 9.1% | 22.2% |

\*The date of symptoms onset was available for only 230 out of 282 donors. Therefore, the proportion of seronegative donors at first donation was calculated using the information from these 230 individuals. The median time after symptoms onset for the first donation was 52 days.

**Table 2:** Characteristics of the 15 repeat COVID convalescent plasma donors.

| Time after symptoms onset<br>and first donation (days) | Time after symptoms onset<br>and last donation (days) | Age<br>Median (range) | Sex |  |
| --- | --- | --- | --- | --- |
|  |  |  | Male (n) | Female<br>(n) |
| 46 (33-77) | 98 (66-114) | 56 (20-67) | 11 | 4 |

As shown in Figure 1A, the level of anti-RBD antibodies at the first donation varies greatly between donors. However, a decrease in anti-RBD antibody level between first and last donation was observed for all donors. To better illustrate the evolution of the anti-RBD antibody response over time, the relative level of anti-RDB antibodies was calculated at each time point using the first time point as reference (Figure 1B). In some donors, an increase was observed after their first donation, but this was always followed by a decline in anti-RBD antibodies at later time points.

**Figure 1.**
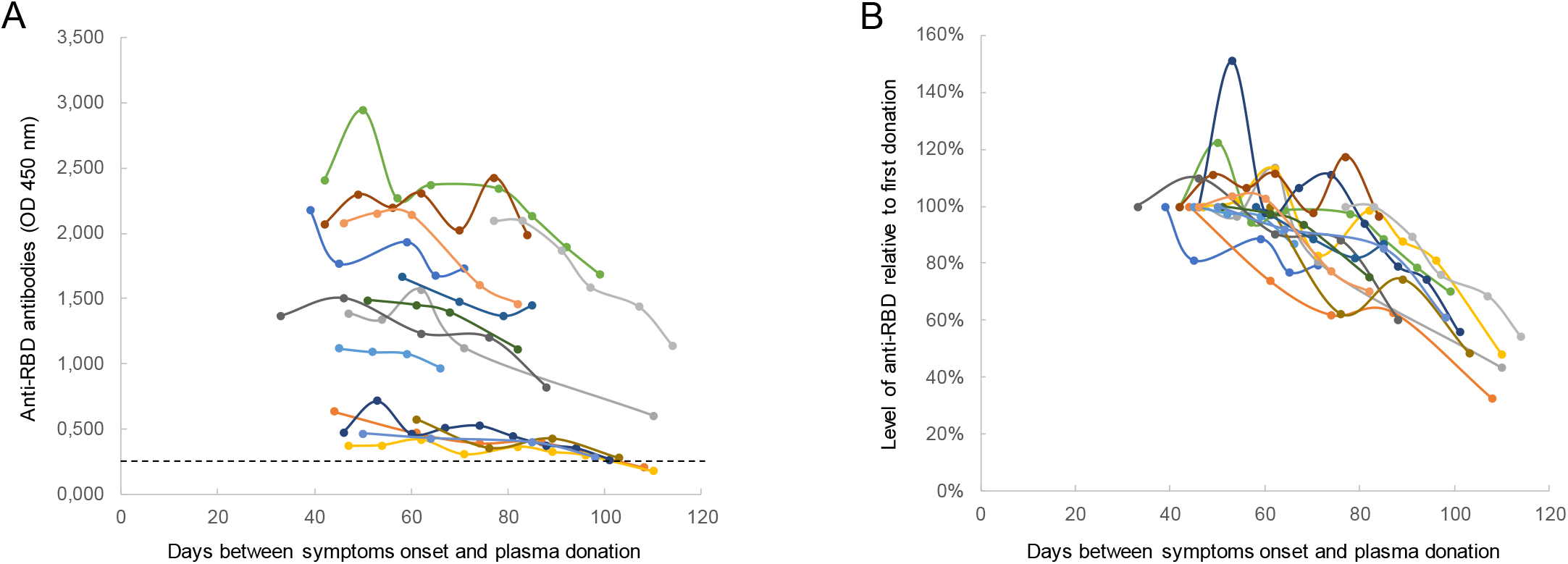
Longitudinal analysis of the anti-RBD antibody response in fifteen repeat CCP donors. The level of anti-RBD antibodies was determined in our semi-quantitative in-house ELISA, using a 1:100 dilution of plasma. (**A**) Each curve represents the OD_450nm_ obtained with the plasma of one donor at every donation (4 to 9 donations per donor) as a function of the days after symptoms onset. The dotted line represents the cut-off value of the ELISA; some donors became seronegative at their last donation. (**B**) Same results but presented as the relative anti-RBD antibody level calculated at each time point using the first time point as reference: 1-[OD_450nm_ at each donation/OD450nm at first donation] x 100.

To rule out the possibility that the decline observed in all donors was a consequence of repeated donations, we determined the correlation between the number of donations and the overall decline in anti-RBD level, as defined using the maximal OD (generally but not always at the first donation) and the OD at the last donation (OD_last donation_/OD_max_). As shown in Figure 2A, the decrease in anti-RBD levels did not correlate with the number of donations (r = 0.127, p-value = 0.6509). We then compared the decrease in anti-RBD level as a function of the time elapsed between the onset of symptoms and the time of the last donation (Figure 2B). The results revealed a significant correlation between these two parameters (r = 0.821, p-value = 0.0002), indicating that the anti-RBD response wanes over time of convalescence rather than because of repeated donations.

**Figure 2.**
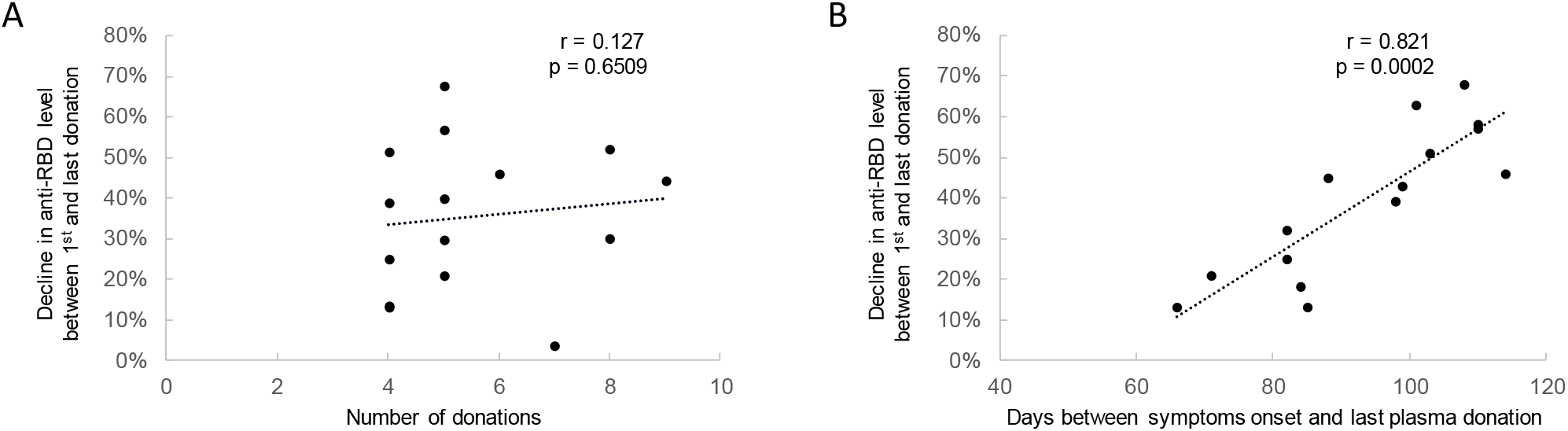
Decline in anti-RBD antibody levels as a function of the number of donations or time elapsed between symptoms onset and last donation. (**A**) Correlation between the number of donations by each donor and the overall decline in anti-RBD level calculated using the following formula: 1-[OD_450nm_ at the last donation/ maximal OD_450nm_ obtained] x 100. (**B**) Correlation between the number of days between symptoms onset and the last donation with the overall decline in anti-RBD level for each donor.

To get a more general picture of the decline in anti-RBD antibodies in CCP donors over time, we performed a repeated measure analysis using a mixed model with participant-level as random effect and time since symptoms onset as fixed effect with adjustment for donor age and sex. For group comparison, the time from onset of symptoms (33-114 days) was divided in quartiles containing similar numbers of samples (from 19 to 22 donor samples) and the data (OD values) in each of these quartiles were combined regardless of the donor identity. Figure 3 shows the distribution, median and mean OD in each quartile. Overall, a significative decrease in OD value from baseline through last donation was observed (p < 0.0001). Pairwise comparisons showed that in the 1^st^ and 2^nd^ quartiles (33 to 53 and 54 to 69 days after symptoms onset respectively), the median and mean OD were quite similar (mean of 1.499 ± 0.760 and 1.309 ± 0.710, median of 1.486 (IQR 1.44) and 1.363 (IQR 1.43), respectively with a p-value of 0.313), although a slight decrease in the mean values could be observed. This suggests that the anti-RBD response is relatively stable during the first months of convalescence. No significant decrease in median and mean OD values was observed between the 2^nd^ and 3^rd^ quartiles (54 to 69 and 70 to 84 days after symptoms onset, respectively) (mean of 1.309 ± 0.710 and 1.321 ± 0.720, median of 1.363 (IQR 1.43) and 1.411 (IQR 1.52), respectively with a p-value of 0.1205). However, the most striking observation comes from the comparison of the 3^rd^ and 4^th^ quartile (70 to 84 and 85 to 114 days after symptoms onset, respectively), where a marked decrease in the mean OD values (significative mean OD decrease of −0.486 from 1.321 ± 0.720 to 0.835 ± 0.670, representing a 36.8% decrease with p-value of 0.0052) and an even more pronounced decrease in median values (median OD decreases from 1.411 (IQR 1.52) to 0.411 (IQR 1.15) representing a 70.1% decrease) were observed. Altogether, these observations are in agreement with recent studies reporting a decrease in anti-RBD responses and neutralization activity in the plasma of convalescent donors a few weeks after symptoms resolution (19–22)

**Figure 3.**
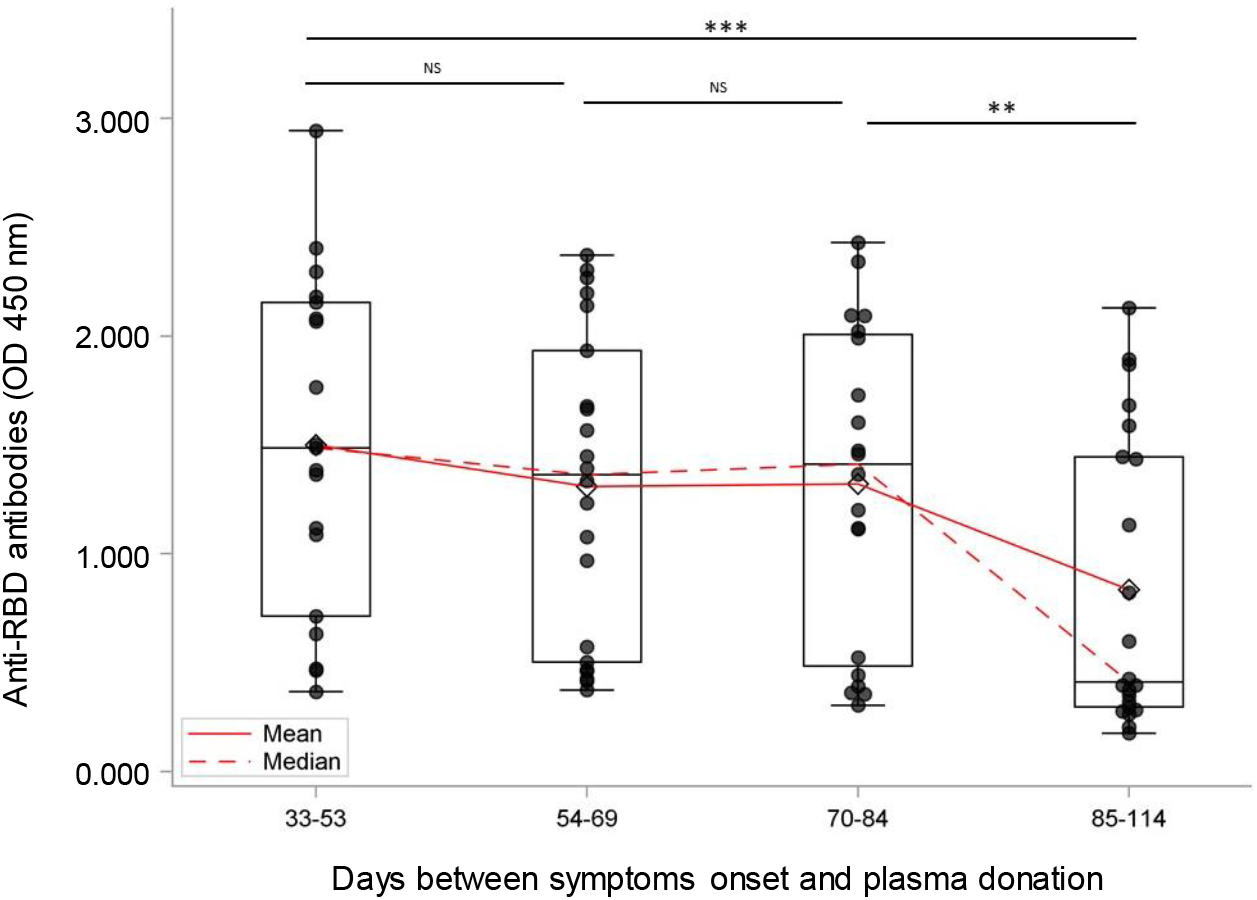
Evolution of the anti-RBD antibody response over time in repeat CCP donors. The time from onset of symptoms (33-114 days) was divided in quartiles containing similar numbers of samples (from 19 to 22 donor samples). The mean and median OD_450nm_ were calculated using all samples in each quartile. Each sample is represented by a dot. Boxes and horizontal bars denote interquartile range (IQR) while horizontal line and lozenge in boxes correspond to median and mean value, respectively. Whisker endpoints are equal to the maximum and minimum values below or above the median ±1.5 times the IQR. Statistical significance was noted as NS, not significant; * p < 0.05; ** p < 0.01; and *** p < 0.001.

Interestingly, the decrease in mean and median OD values during a period of about 20 days (considering the mean and median of 3^rd^ and 4^th^ quartiles, both of 76 and 95 days respectively) is reminiscent of the plasma IgG half-life of 21 days (23), suggesting that *de novo* synthesis of anti-RBD antibodies stopped between the 3^rd^ and 4^th^ quartiles in all CCP donors. This time frame is consistent with the first wave of a humoral immune response during which short-lived plasma cells actively secrete pathogen-specific antibodies until the antigen is eliminated (24). This is usually followed by the emergence of a cellular memory response that could play a major role in the long-term protection against reinfection, as recently proposed (25).

Our study contains some limitations as only anti-RBD antibodies were measured. Additional work including the characterization of our CCP donor plasma samples on other SARS-CoV-2 antigens (eg. full spike, nucleocapsid) will permit to extend our initial observations on RBD antibodies to a broader humoral response to SARS-CoV-2 and help to better define its persistence. Nevertheless, the availability of sequential samples from the CCP repeated donors permitted to better pinpoint the time at which the anti-RBD response starts to significantly decline, regardless of the initial anti-RBD antibody level which has been shown to correlate with disease severity (19,26,27). Consequently, individuals who experienced a mild or asymptomatic COVID-19 could become seronegative within a period as short as 100 days after disease onset. This observation has important implications for seroprevalence studies in the general population. Such studies should be performed close to the peak of infection, when most infected individuals (symptomatic or not) will still have easily detectable SARS-CoV-2 antibodies, to better estimate the true number of SARS-CoV-2 infections.

## METHODS

### Ethics statement

All work was conducted in accordance with the Declaration of Helsinki in terms of informed consent and approval by an appropriate institutional board. Convalescent plasmas were obtained from donors who consented to participate in this research project at Héma-Québec (REB # 2020-004).

### Convalescent plasma donors

Recovered COVID-19 patients were recruited mostly following self-identification and through social media. All participants have received a diagnosis of COVID-19 by the Québec Provincial Health Authority and met the donor selection criteria for plasma donation in use at Héma-Québec. They were allowed to donate plasma at least 14 days after complete resolution of COVID-19 symptoms. Males and females with no history of pregnancy meeting the above criteria were invited to donate plasma, after informed consent. A volume of 500 mL to 750 mL of plasma was collected by plasmapheresis (TRIMA Accel®, Terumo BCT). Seropositive donors were allowed to donate additional plasma units every six days, for a maximum of 12 weeks. Disease severity (date of symptoms onset, end of symptoms, type and intensity of symptoms, need for hospitalization/ICU) was documented for each donor using a questionnaire administered at the time of recruitment.

### SARS-CoV-2 RBD ELISA

The presence of antibodies against SARS-CoV-2 RBD was determined using a semi-quantitative ELISA. The assay was adapted from a recently described protocol (20,21). The plasmid encoding for SARS-CoV-2 S RBD was synthesized commercially (Genscript). The RBD sequence (encoding for residues 319-541) fused to a C-terminal hexahistidine tag was cloned into the pcDNA3.1(+) expression vector. Recombinant RBD proteins were produced in transfected FreeStyle 293F cells (Invitrogen) and purified by nickel affinity chromatography. Recombinant RBD was diluted to 2.5 μg/mL in PBS (Fisher Scientific) and 100 μl of the dilution was distributed in the wells of flat-bottom 96-well microplates (Immulon 2HB; Thermo Scientific). The plates were placed overnight at 2-8°C for antigen adsorption. During validation, we determined that the plates should be used within 48 hours of antigen adsorption. For the assay, the plates were emptied and a volume of 300 μl/well of blocking buffer (PBS-0.1% Tween (Sigma)-2% BSA (Sigma)) was added. The microplates were incubated for one hour at room temperature (RT) followed by washing four times (ELx405 microplate washer, Bio-Tek) with 300 μL/well of washing solution (PBS-0.1% Tween). Plasma samples were diluted 1:100 in blocking buffer and a volume of 100 μl of each diluted sample was added to the microplate wells in triplicate. The human monoclonal anti-SARS-CoV CR3022, known to cross-react with SARS-CoV-2 RBD (28), was included in each microplate at 50 ng/mL and served as positive control. Commercial plasma (SeraCon, SeraCare Life Sciences, Inc.) prepared from pools of 1 000 liters of plasma collected before SARS-CoV-2 outbreak was used at a 1:100 dilution as negative control. The plates were incubated for 1 hour at RT followed by washing and addition of 100 μl of anti-human IgA + IgG + IgM (H+L) conjugated to HRP (Jackson ImmunoResearch Laboratories, Inc.) diluted 1:50 000 in blocking buffer. The plates were incubated once again for one hour at RT followed by washing and addition of 100 μl of 3,3’,5,5’-Tetramethylbenzidine (TMB, ESBE Scientific). The colorimetric reaction proceeded for 20 minutes at RT and was stopped by addition of 100 μl of H2SO4 1N (Fisher Scientific). The plates were then read within 30 minutes at 450 nm using a Synergy H1 microplate reader (Bio-Tek). The OD cut-off for seropositivity was set at 0.250.

### Statistical analysis

Changes from baseline measurements were modeled with the use of a linear mixed-effects model for repeated measures based on a participant-level analysis with fixed effects for sex, age and time since symptoms onset. Compound symmetry covariance matrix was used to model the within-patient variance–covariance errors with degrees of freedom controlled using a Satterthwaite approximation to account for inexact F distributions of the fixed effects in repeated-measure mixed model (29,30). Pairwise post-hoc tests were estimated using Tukey Kramer adjustment for multiple comparisons.

## Acknowledgments

The authors are grateful to the convalescent plasma donors who participated in this study and the Héma-Québec team involved in convalescent donor recruitment and plasma collection. We also thank Dr M. Gordon Joyce (U.S. MHRP) for the monoclonal antibody CR3022. This work was supported in part by le Ministère de l’Économie et de l’Innovation du Québec, Programme de soutien aux organismes de recherche et d’innovation to A.F. A.F. is the recipient of a Canada Research Chair on Retroviral Entry # RCHS0235 950-232424. G.B.B. and J.P. are supported by CIHR fellowships. P.B. is supported by a FRQS Junior 2 salary award. The funders had no role in study design, data collection and analysis, decision to publish, or preparation of the manuscript. The authors declare no competing interests.

## Notes

### Competing Interest Statement

The authors have declared no competing interest.

## References

1. Duan K, Liu B, Li C, Zhang H, Yu T, Qu J, et al. Effectiveness of convalescent plasma therapy in severe COVID-19 patients. Proc Natl Acad Sci. 2020 Apr 6;202004168.

2. Shen C, Wang Z, Zhao F, Yang Y, Li J, Yuan J, et al. Treatment of 5 Critically Ill Patients With COVID-19 With Convalescent Plasma. JAMA. 2020 Apr 28;323(16):1582.

3. Zhang B, Liu S, Tan T, Huang W, Dong Y, Chen L, et al. Treatment with convalescent plasma for critically ill patients with SARS-CoV-2 infection. Chest. 2020 Mar;S0012369220305717.

4. Casadevall A, Pirofski L. The convalescent sera option for containing COVID-19. J Clin Invest. 2020 Mar 13;10.1172/JCI138003.

5. Dzik S. COVID-19 Convalescent Plasma: Now Is the Time for Better Science. Transfus Med Rev. 2020 Apr;S0887796320300262.

6. Ye M, Fu D, Ren Y, Wang F, Wang D, Zhang F, et al. Treatment with convalescent plasma for COVID-19 patients in Wuhan, China. J Med Virol [Internet]. 2020 Apr 15 [cited 2020 May 13]; Available from: http://doi.wiley.com/10.1002/jmv.25882

7. Zeng Q-L, Yu Z-J, Gou J-J, Li G-M, Ma S-H, Zhang G-F, et al. Effect of Convalescent Plasma Therapy on Viral Shedding and Survival in COVID-19 Patients. J Infect Dis. 2020 Apr 29;jiaa228.

8. Rojas M, Rodríguez Y, Monsalve DM, Acosta-Ampudia Y, Camacho B, Gallo JE, et al. Convalescent plasma in Covid-19: Possible mechanisms of action. Autoimmun Rev. 2020 Jul;19(7):102554.

9. Rajendran K, Krishnasamy N, Rangarajan J, Rathinam J, Natarajan M, Ramachandran A. Convalescent plasma transfusion for the treatment of COVID-19: Systematic review. J Med Virol 2020 May 12;jmv.25961.

10. Gharbharan A, Jordans CCE, Geurtsvankessel C. Convalescent Plasma for COVID-19. A randomized clinical trial.:16.

11. Li L, Zhang W, Hu Y, Tong X, Zheng S, Yang J, et al. Effect of Convalescent Plasma Therapy on Time to Clinical Improvement in Patients With Severe and Life-threatening COVID-19: A Randomized Clinical Trial. JAMA [Internet]. 2020 Jun 3 [cited 2020 Jun 4]; Available from: https://jamanetwork.com/journals/jama/fullarticle/2766943

12. Robbiani DF, Gaebler C, Muecksch F, Cetrulo Lorenzi J, Wang Z, Cho A, et al. Convergent Antibody Responses to SARS-CoV-2 Infection in Convalescent Individuals [Internet]. Immunology; 2020 May [cited 2020 May 19]. Available from: http://biorxiv.org/lookup/doi/10.1101/2020.05.13.092619

13. Ni L, Ye F, Cheng M-L, Feng Y, Deng Y-Q, Zhao H, et al. Detection of SARS-CoV-2-specific humoral and cellular immunity in COVID-19 convalescent individuals. Immunity. 2020 May;S1074761320301813.

14. Wu F, Wang A, Liu M, Wang Q, Chen J, Xia S, et al. Neutralizing antibody responses to SARS-CoV-2 in a COVID-19 recovered patient cohort and their implications [Internet]. Infectious Diseases (except HIV/AIDS); 2020 Apr [cited 2020 Apr 9]. Available from: http://medrxiv.org/lookup/doi/10.1101/2020.03.30.20047365

15. Jackson LA, Anderson EJ, Rouphael NG, Roberts PC, Makhene M, Coler RN, et al. An mRNA Vaccine against SARS-CoV-2 — Preliminary Report. N Engl J Med [Internet]. 2020 Jul 14 [cited 2020 Jul 16]; Available from: https://doi.org/10.1056/NEJMoa2022483

16. Zhang G, Nie S, Zhang Z, Zhang Z. Longitudinal Change of Severe Acute Respiratory Syndrome Coronavirus 2 Antibodies in Patients with Coronavirus Disease 2019. J Infect Dis. 2020 Jun 29;222(2):183–8.

17. Long Q-X, Liu B-Z, Deng H-J, Wu G-C, Deng K, Chen Y-K, et al. Antibody responses to SARS-CoV-2 in patients with COVID-19. Nat Med [Internet]. 2020 Apr 29 [cited 2020 May 1]; Available from: http://www.nature.com/articles/s41591-020-0897-1

18. Lynch KL, Whitman JD, Lacanienta NP, Beckerdite EW, Kastner SA, Shy BR, et al. Magnitude and kinetics of anti-SARS-CoV-2 antibody responses and their relationship to disease severity [Internet]. Infectious Diseases (except HIV/AIDS); 2020 Jun [cited 2020 Jul 1]. Available from: http://medrxiv.org/lookup/doi/10.1101/2020.06.03.20121525

19. Long Q-X, Tang X-J, Shi Q-L, Li Q, Deng H-J, Yuan J, et al. Clinical and immunological assessment of asymptomatic SARS-CoV-2 infections. Nat Med [Internet]. 2020 Jun 18 [cited 2020 Jul 10]; Available from: http://www.nature.com/articles/s41591-020-0965-6

20. Beaudoin-Bussières G, Laumaea A, Anand SP, Prévost J, Gasser R, Goyette G, et al. Decline of humoral responses against SARS-CoV-2 Spike in convalescent individuals. bioRxiv. 2020 Jul 15;2020.07.09.194639.

21. Prévost J, Gasser R, Beaudoin-Bussières G, Richard J, Duerr R, Laumaea A, et al. Cross-sectional evaluation of humoral responses against SARS-CoV-2 Spike. BioRxiv Prepr Serv Biol. 2020 Jun 10;2020.06.08.140244.

22. Seow J, Graham C, Merrick B, Acors S, Steel KJA, Hemmings O, et al. Longitudinal evaluation and decline of antibody responses in SARS-CoV-2 infection [Internet]. Infectious Diseases (except HIV/AIDS); 2020 Jul [cited 2020 Jul 16]. Available from: http://medrxiv.org/lookup/doi/10.1101/2020.07.09.20148429

23. Morell A, Terry WD, Waldmann TA. Metabolic properties of IgG subclasses in man. J Clin Invest. 1970 Apr 1;49(4):673–80.

24. Amanna IJ, Slifka MK. Mechanisms that determine plasma cell lifespan and the duration of humoral immunity: Long-term antibody production. Immunol Rev. 2010 Jun 15;236(1):125–38.

25. Sekine T, Perez-Potti A, Rivera-Ballesteros O, Strålin K, Gorin J-B, Olsson A, et al. Robust T cell immunity in convalescent individuals with asymptomatic or mild COVID-19 [Internet]. Immunology; 2020 Jun [cited 2020 Jul 12]. Available from: http://biorxiv.org/lookup/doi/10.1101/2020.06.29.174888

26. Woloshin S, Patel N, Kesselheim AS. False Negative Tests for SARS-CoV-2 Infection — Challenges and Implications. N Engl J Med. 2020 Jun 5;NEJMp2015897.

27. Mallapaty S. WILL CORONAVIRUS ANTIBODY TESTS REALLY CHANGE EVERYTHING? Nature. 2020;580:571–2.

28. ter Meulen J, van den Brink EN, Poon LLM, Marissen WE, Leung CSW, Cox F, et al. Human Monoclonal Antibody Combination against SARS Coronavirus: Synergy and Coverage of Escape Mutants. Burton DR, editor. PLoS Med. 2006 Jul 4;3(7):e237.

29. Satterthwaite FE. An Approximate Distribution of Estimates of Variance Components. Biom Bull. 1946;2(6):110–4.

30. Laird NM, Ware JH. Random-Effects Models for Longitudinal Data. Biometrics. 1982;38(4):963–74.

